# The PDE4D ortholog Dunce suppresses memory in *Drosophila melanogaster*

**DOI:** 10.64898/2025.11.29.691281

**Authors:** Timo Hasselmann, Marcel Verbrüggen, Marie Müller, Magdalena Gompert, Henrike Scholz

## Abstract

The regulation of cAMP concentration is a key component of the mechanisms underlying learning and memory. In *Drosophila melanogaster*, the cAMP-specific phosphodiesterase mutant *dunce^1^* results in increased levels of cAMP but poorer associative short-term memory and anesthesia-resistant memory. This raises the question of how elevated cAMP levels can cause defects in both short-term and long-term memory. To answer this question, we analyzed associative olfactory learning and memory in *Drosophila* larvae of both sexes with the isoform-specific *dunce^Δ143^* mutation. Flies with this mutation exhibited better learning than flies with the *dunce^1^* mutant. The memory facilitated by *dunce^Δ143^* was stable and easily reversable. The improvement in memory was also traced to a defined subset of neurons. A comparison of the subcellular localization of Dunce isoforms indicates that the regulation of cAMP in the soma suppresses early memory formation. We revealed that Dunce regulates at least two different aspects of learning and memory, specifically, preventing the premature formation of memory and facilitating the formation of memory after multiple training sessions. The different functions of Dunce might be due to different cell types.

**Significance Statement:** In model systems such as Aplysia and mice, elevated cAMP levels are associated with increased learning and memory performance; in contrast, *Drosophila dunce* mutants with elevated cAMP levels exhibit memory decline. Here, we revisited the function of the phosphodiesterase 4D ortholog Dunce and showed that Dunce both promotes and inhibits memory formation.

## Introduction

In various organisms, including invertebrates such as *Drosophila melanogaster*, the second messenger cyclic adenosine monophosphate (cAMP) is crucial for the regulation of neuronal function. cAMP regulates neuronal excitability, synaptic plasticity, and learning and memory (Argyrousi et al., 2020; Lee, 2015). Neurons are highly differentiated cells with dendrites, soma, axons and axon terminals. At the pre-synapse, cAMP levels regulate synaptic neurotransmitter release, whereas in the postsynaptic region, cAMP affects neuronal excitability (Fukaya et al., 2023; Argyrousi et al., 2020; Kidokoro et al., 2004; Lee, 2015).

To ensure that cAMP can regulate different processes in neurons, cAMP levels must be tightly controlled in both space and time. cAMP-dependent phosphodiesterases (PDEs) are essential regulators of cAMP signaling. These enzymes hydrolyze cAMP to 5’AMP and thereby terminate the action of cAMP (Conti and Beavo, 2007). The specific localization of different PDEs creates subcellular cAMP microdomains, restricting signaling to designated areas and preventing cAMP diffusion (Houslay, 2010).

At the level of behavior, in mice, blocking PDE activity in the hippocampus improves memory, supporting the role of cAMP elevation in memory formation and enhancement (Peters et al., 2014). In contrast, the *Drosophila* PDE4D mutant *dunce^1^* (*dnc^1^*), which exhibits elevated cAMP levels, was originally identified due to deficits in olfactory learning and memory (Dudai et al., 1976; Lee, 2015). The *dnc^1^*mutant shows reduced short-term memory due to defects in anesthesia-resistant memory. These defects can be attributed to dysregulation of PDE activity in the local interneurons of the antennal lobes and neurons of the mushroom bodies, a structure crucial for learning and memory in *Drosophila* (Scheunemann et al., 2012). In addition to these defects in short-term memory, *dnc^1^* shows improvements in long-term memory, similar to what is observed when Dnc expression is reduced in serotonergic SPN neurons (Scheunemann et al., 2012). At the cellular level, the application of different neurotransmitters to mushroom bodies triggers increases in cAMP levels in various subcellular compartments. However, these changes are absent in *Drosophila*,*dnc* mutants which lack the cAMP-specific phosphodiesterase (Gervasi et al., 2010; Day et al., 2005; Davis and Kiger, 1981).

In the developing nervous system of *Drosophila* larva, *dunce* is also involved in memory regulation. *dnc^1^* mutant larvae show learning and memory deficits in olfactory associative short-term memory, similar to what is observed in adult flies (Aceves-Pina and Quinn, 1979; Khurana et al., 2009; Honjo and Furukubo-Tokunaga, 2009; Honjo and Furukubo-Tokunaga, 2005). However, unlike *dnc* mutant adults, *dnc* mutant larvae form anesthesia-resistant memory (Widmann et al., 2016). Distinct subcellular compartments of cAMP have been identified in motor neurons at the neuromuscular junction of *Drosophila* larvae. These neurons exhibit three independent cAMP signaling compartments, located in the soma, the axon, and the axon terminal. However, blocking PDE activity leads to a uniform distribution of an Epac1-based cAMP sensor in these cells (Maiellaro et al., 2016).

Several promotors regulate *Dunce* transcription, and at least eight different *Dunce* transcripts are generated through alternative splicing (Qiu and Davis, 1993; Ruppert et al., 2017). These transcripts are categorized into five classes of isoforms according to their unique N-terminal regions and the presence of the UCR1 and UCR2 domains in addition to the catalytic PDE domain (Ruppert et al., 2017). Different isoforms are expressed in different subcellular compartments (Ruppert et al., 2017). In *dnc^1^* mutants, several different transcripts are dysregulated (downregulated or upregulated) (Ruppert et al., 2017), making it difficult to attribute the defects in learning and memory in *dnc^1^* to a specific Dunce isoform.

Here, we characterize the learning and memory abilities of larvae with the isoform-specific *dnc^Δ143^* mutation using an associative olfactory learning and memory paradigm. Unlike *dnc^1^* mutants, the *dnc^Δ143^*mutants lack a specific isoform, Dnc^PA^ (Ruppert et al., 2017). Using a *dnc^RA^* promotor Gal4 driver to control the expression of different Dnc isoforms, we identified the neurons in which PDE activity is required to regulate learning and memory. Moreover, we analyzed the expression of different Dnc isoforms in motor neurons to determine where Dnc is required at the cellular level. Our data provide evidence that *Dunce* mutants with elevated cAMP levels are not “dunces” at all but in fact can learn better than wild-type *Drosophila*.

## Materials and Methods

### Drosophila stocks

Flies were maintained on standard cornmeal-based food at 25°C and 60% relative humidity on a 12 h light/dark cycle. For the behavioral experiments, the flies were reared under density control; e.g., 30 virgin flies were crossed with 15 male flies. All experiments were performed with foraging three- to five-day-old 3^rd^-instar larvae. The following stocks were previously generated in our laboratory in the *w^1118^* background from the Scholz laboratory: *w^1118^,dnc*^Δ*143*^; *w^1118^;UAS-dnc^RA^::eGFP, w^1118^;UAS-eGFP-dnc^RA^*, *w^1118^;UAS-dnc^RG^::eGFP, w^1118^;dnc^RA^-Gal4 w^1118^,dnc*^Δ*143*^*;dnc^RA^-*Gal4; *w^1118^,dnc*^Δ*143*^*;UAS-dnc^PDE^*; *w^1118^,dnc*^Δ*143*^*;UAS-eGFP-dnc^RA;^ w^1118^,dnc*^Δ*143*^*; UAS-dnc^RA^::eGFP; w^1118^,dnc*^Δ*143*^*;UAS-dnc^RG^::eGFP* (Ruppert et al., 2017). The *CantonS* BDSC_64349, *dnc^1^* BDSC_6020, and *UAS-mCD8::GFP* BDSC_108068 stocks were obtained from the Bloomington *Drosophila* Stock Center. *UAS-dnc^KK-107967^* (Fenckova et al., 2019) was obtained from the Schenck laboratory, and *DvGlut-Gal4* (Collins et al., 2006) was obtained from the Collins laboratory.

### Associative olfactory learning and memory

The assay was conducted using 2 M NaCl as a negative reinforcer as described by (Widmann et al., 2016). Briefly, a group of 20 larvae were trained for 5 min on 2.5% agarose plates containing 2 M NaCl in the presence of odorant A followed by a five min exposure to odorant B on a second agarose plate. A second group of larvae was reciprocally trained (exposed to odorant B first) on a reinforcer-containing agarose plate. The training was performed once or repeated three times. To test whether the larvae formed an association between the odorant and the reward, the flies were given the choice between odorant A and B on the reinforcer containing the agarose plate at 2 min after the training. The alterations from the procedure are indicated in the text. Benzaldehyde (Sigma–Aldrich #12010), ethyl acetate (1:50; Sigma–Aldrich # 58958) and amyl acetate (1:100; Emplura #8.18700) were used as odorants. To analyze whether odorant perception or reward processing is altered in the larvae, preference was determined on agarose plates with and without reinforcers. In addition, an odorant balance test was performed to determine whether the animals perceived the odorants as equally rewarding—prerequisites for the learning and memory experiments.

To analyze odorant preference, a group of 20 larvae were placed in the neutral zone, a 1-cm-wide area in the center of the plate. They were allowed to choose between cups filled with the respective odorants and a cup filled with the solvent paraffin. To analyze whether both odorants were perceived and evaluated equally, two different odorants were placed on opposite sides of the agarose plate. The animals were allowed 5 min to choose before the number of larvae was counted. The preference index was calculated as the number of larvae that moved toward the odorant, divided by the total number of larvae. When two odorants were present in the test situation, the preference index was calculated as the number of larvae that moved away for the reinforced odorant minus the number of larvae that moved toward the nonreinforced odorant, divided by the total number of larvae. To calculate the learning index, the preference indices of the reciprocally trained groups of larvae were summed and divided by two.

### Immunohistochemistry

The CNSs of late 3^rd^-instar larvae were dissected in ice-cold PBS and fixed with 3.7% formaldehyde/PBS for 20 min. After three rinses, the samples were washed three times for 20 min in PBS supplemented with 0.3% Triton (PBT) and then blocked with 5% FCS/PBT for 2 h. The samples were incubated with the primary antibody in 5% FCS/PBT on a shaker at 4°C for two nights. The samples were rinsed and washed again for at least 1 h and incubated with the secondary antibody in 0.3% PBT at 4°C overnight. Then, the CNSs were again rinsed and washed again for 1 h, incubated in 50% glycerol/PBS for 30 min and mounted.

To analyze the NMJ, wandering late 3^rd^-instar larvae were placed on ice, and the body wall, including the CNS, was dissected in ice-cold PBS. Multiple genotypes were pinned to the same dissection plate and not removed until mounting to ensure equal treatment. The samples were fixed in 3.7% formaldehyde/PBS for 15 min. The samples were then rinsed, washed for 1 h with PBS supplemented with 0.1% Triton, and blocked in 5% FCS/PBT for 30 min. The tissue was incubated with the primary antibody in 5% FCS/PBT/or PBS at 4°C overnight. After being rinsed and washed, the samples were incubated with the secondary antibody in 0.3% PBT for 2 h at room temperature. The samples were then rinsed and washed for 1 h before embedding. The samples were mounted in VECTASHIELD ^®^ and imaged with an Olympus Fluoview FV1000 or a Leica SP8. The image stacks were analyzed using Fiji ImageJ and processed with Photoshop CS6. At the NMJ, segment A4 at NMJ6/7 was analyzed.

The following antibodies were used: anti-GFP rabbit (1:1000, Rockland Immunochemicals #600-401-215), anti-nc82 mouse (1:20), anti-FasII mouse (1:50), anti-DLG mouse (1:1000), anti-ChAT mouse (1:100), anti-Lamin mouse (1:100), Cy3-conjugated anti-HRP goat (1:500, Jackson Immuno-Research Ltd. #123-165-021), anti-rabbit AF488 (1:500, Invitrogen #A10522), anti-mouse AF633 (1:500, Invitrogen #A-21050) and anti-mouse Cy3 (1:500, Jackson ImmunoResearch #115-165-146).

### Predictive models for Dnc proteins

The structures of the Dnc^PA^ fusion proteins were analyzed with AlphaFold2 (Jumper et al., 2021) using the pipeline and parameters published in the AlphaFold Google Collaboratory notebook with default settings except that the relaxation step was disabled. The resulting models were analyzed and rotated using PyMOL2.6 and visualized using PhotoshopC6.

### Data analysis and statistics

To determine whether the data were normally distributed, the Shapiro–Wilk test (significance level *P* < 0.05) was used. One-sample t tests were used to calculate differences from random choice using R (R Core Team, 2021) and https://www.statskingdom.com/. For normally distributed data, significance was determined by one-way ANOVA combined with the Tukey–Kramer *post hoc* HSD test.

For experiments with only two genotypes, Student’s t test was used; for comparisons among more than two groups, ANOVA with Tukey’s honest significant difference (HSD) test was used. In the figures, the behavioral data are presented as box plots that include the data points. In the figure legends, the data are presented as the means ± s.e.m.s. In addition to being shown in the figures, the data, including statistics, are provided in Table S1.

## Results

Larvae can learn to associate an odorant with a negative reward, such as an electric shock or 2 M NaCl. Depending on the training regimen, they remember to avoid the punished odorant in the short term, middle term or long term (Khurana et al., 2009). *dnc1*-mutant larvae exhibit defects in short-term associative learning and memory, specifically in the anesthesia-sensitive component of memory (Aceves-Pina and Quinn, 1979; Khurana et al., 2009; Widmann et al., 2016). Mutation of *dnc^1^* affects multiple *dunce* transcripts whose expression is downregulated or upregulated (Ruppert et al., 2017). To reevaluate the role of specific isoforms in learning and memory, we wanted to analyze the learning and memory performance of *dnc^Δ143^* with a deficiency in the promoter region of the *dnc^RA^* transcript (Figure 1A; Ruppert et al., 2017).

**Figure 1:**
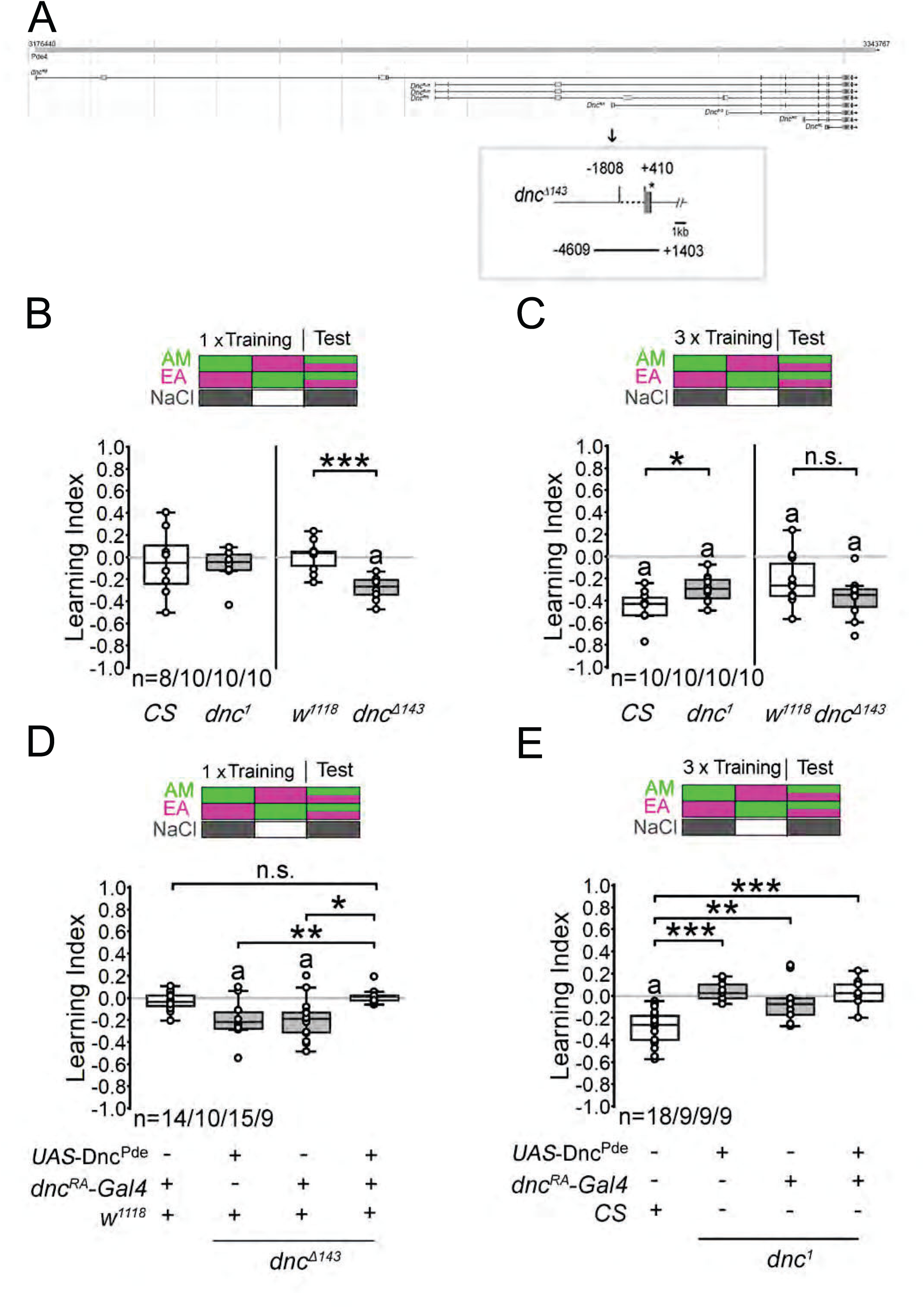
Dnc suppresses learning and memory. ***A*,** Genomic organization of *dunce*. The box shows a higher magnification of the *dnc^RA^* promoter region of *dnc*^Δ*143*^ mutants, and the line below indicates the fragment used to generate *dnc^RA^-Gal4. **B***, After one training session, control animals and *dnc^1^* mutants did not form memories, but *dnc*^Δ*143*^ larvae learned to associate the odorant with 2 M NaCl. Student’s t test: *CS* = -0.05 ± 0.1, *dnc^1^* = -0.07 ± 0.05, Df (16), t = 0.161, *P* = 0.874 and *w^1118^* = 0.01 ± 0.04*, w^1118^dnc*^Δ*143*^ = -0.28 ± 0.03, Df (18); t = -5.210; *P* < 0.0001. ***C***, After three cycles of training, *dnc^1^*learned and remembered significantly less than *CS* larvae, whereas the *dnc*^Δ*143*^ larvae learned and remembered to a similar extent as their respective controls. *CS* = -0.46 ± 0.04, *dnc^1^* = -0.29 ± 0.04, Df (18), t = -2.758, *P* = 0.013 and *w^1118^* = -0.22 ± 0.07*, w^1118^dnc*^Δ*143*^ = -0.37 ± 0.06, Df (18), t = -1.634, *P* = 0.1197. Normally distributed data in ***B*** and ***C*** were analyzed with an unpaired Student’s t test. ***D***, Expression of the Dnc^PDE^ domain under the control of the *dnc^RA^*-Gal4 driver reduced the learning and memory of *dnc*^Δ*143*^ to control levels. ANOVA with Tukey’s HSD: Df (3,47), F = 6.7045; *P* = 0.0008 and *w^1118^; dnc^RA^-*Gal4 = -0.03 ± 0.02; *w^1118^,dnc*^Δ*143*^*;dnc^RA^-*Gal4 = -0.2 ± 0.05; *w^1118^,dnc*^Δ*143*^*;UAS-dnc^PDE^ =* -0.2 ± 0.06; *w^1118^,dnc*^Δ*143*^*; UAS-dnc^PDE^; dnc^RA^-*Gal4 *=* 0.02 ± 0.02. ***E***, The expression did not improve learning or memory after three cycles of training in *dnc^1^* larvae. ANOVA with Tukey’s HSD: Df (3,44), F = 13.6884; *P* < 0.0001 and *CS* = -0.29 ± 0.04; *dnc^1^;dnc^RA^-dnc^RA^-*Gal4 = -0.05 ± 0.06; *dnc^1^;UAS-dnc^PDE^ =* 0.04 ± 0.03; *dnc^1^; UAS-dnc^PDE^; dnc^RA^-dnc^RA^-*Gal4 *=* 0.01 ± 0.05. Significant differences from random choice were determined using a one-sample t test (*P* < 0.05) and indicated with “a”. “n” indicates the number of reciprocally trained independent groups of larvae.

### *dnc^Δ143^* larvae learn better than control larvae

To analyze the learning and memory performance of *dnc^Δ143^* mutant larvae, we analyzed the ability of 3^rd^-instar larvae to learn and remember the association between ethyl acetate (EA; 1:50) or amyl acetate (AM; 1:100) and the negative reinforcer of 2 M NaCl (Figure 1). For comparison, we also included the *dnc^1^* mutants with their respective control *CantonS* in the analysis.

First, to avoid bias in the evaluation of the reinforcer, we tested whether larvae perceived EA or AM similarly on agarose plates or on plates containing 2 M NaCl and whether both odorants were equally attractive (Table S1). Under test conditions, the controls and mutants perceived both odorants as equally attractive. During training, the larvae were exposed to EA or AM on an agarose plate with 2 M NaCl for 5 min. This was followed by a 5 min exposure to a second odorant without reinforcement. During the test, the larvae were allowed 5 min to choose between the two odorants. After one training session, the *dnc^Δ143^* mutants avoided the punished odorant, whereas the *w^1118^*, *CantonS* and *dnc^1^* larvae did not (Figure 1A).

Increasing the number of training sessions increases the stability of memory in the larvae (Khurana et al., 2009). To investigate the effect of additional training on memory, we analyzed memory formation after three training cycles in *dnc^Δ143^* mutants and *dnc^1^*mutants (Figure 1B).

Here, we were able to replicate previous results, namely, that *dnc^1^*memory was approximately 36% worse than that of the control *CantonS* (Figure 1B; Khurana et al., 2009). In contrast, *dnc*^Δ*143*^ mutants learned and remembered just as well as the respective controls (Figure 1B). To analyze whether d*nc*^Δ*143*^ mutants learn and remember different odorant pairs more effectively after one training cycle, we replicated the experiments using a different combination of odorants: benzaldehyde (BA) and AM (Figure S1). Once again, the *dnc*^Δ*143*^ mutant larvae learned better after one training cycle, but after further training, their learning was indistinguishable from that of the control larvae (Figure S1 A and S1B). After one training cycle, *CantonS* larvae remembered the punished odorant, whereas *dnc^1^* showed a significant reduction in learning and memory, consistent with previous results (Widmann et al., 2016; Figure S1A). Similar learning and memory performance were observed in *CantonS* and *dnc^1^* larvae when the number of training cycles was increased to three. The learning and memory defects of *dnc^1^* mutants appeared to be influenced by the odorant pair. In contrast, the improved learning and memory of *dnc*^Δ*143*^ did not appear to be specific to the odorant pair.

Elevated cAMP levels are observed in *dnc*^Δ*143*^ mutants (Ruppert et al., 2017). To determine whether reducing cAMP levels could repress learning and memory deficits, we expressed PDE using the UAS-*dnc^PDE^* transgene under the control of the *dnc^RA^*-Gal4 driver in *dnc*^Δ*143*^ mutants. We then analyzed learning performance after a single training cycle (Figure 1C). The *UAS-dnc^PDE^* transgene comprises mainly the PDE domain, which is present in all Dnc isoforms and efficiently reduces cAMP levels (Cheung et al., 1999). The expression of the Dnc PDE domain reduced enhanced learning and memory in *dnc*^Δ*143*^ mutants to control levels (Figure 1C). This led us to two conclusions, namely, that the Dnc PDE is necessary to suppress memory after a single training cycle and that the neurons in which PDE activity is required are targeted by the *dnc^RA^*-Gal4 driver.

Similar to *dnc*^Δ*143*^ mutants, *dnc^1^* mutants exhibit increased cAMP levels (Byers et al., 1981). However, expression of the Dnc^PDE^ domain did not improve the memory performance of *dnc^1^*-positive cells after three training cycles (Figure 1D). Therefore, the elevated cAMP levels in *dnc^RA^*-positive neurons cannot be the cause of the learning and memory deficits of *dnc^1^*-mutant larvae.

### The aversive memory formed by *dnc***^Δ^***^143^* is stable but flexible

To investigate which type of learning and/or memory is enhanced in *dnc*^Δ*143*^ mutants, we first analyzed how stable the memory is (Figure 2A). We determined that the improved memory of the *dnc*^Δ*143*^ mutants was maintained for up to three hours after training. Next, we determined whether the memory was anesthesia resistant by administering a cold shock immediately after training and measuring learning and memory 10 minutes later (Figure 2B). The individuals that received a cold shock immediately after training did not exhibit any detectable learning and/or memory, indicating that the memory formed by the larvae directly after training is anesthesia sensitive, a characteristic that is shared with the formation of long-term memory. To further distinguish the memory defects observed in *dnc^1^* and *dnc*^Δ*143*^, we analyzed the stability of the memory of *dnc^1^*mutants after three cycles of learning (Figure 2C). Consistent with previous results after three cycles of training, *CantonS* larvae develop a memory that declines but remains measurable over a period of 180 min (Figure 2C and (Widmann et al., 2016). However, the memory of *dnc^1^* mutants disappeared within 30 min after training.

**Figure 2:**
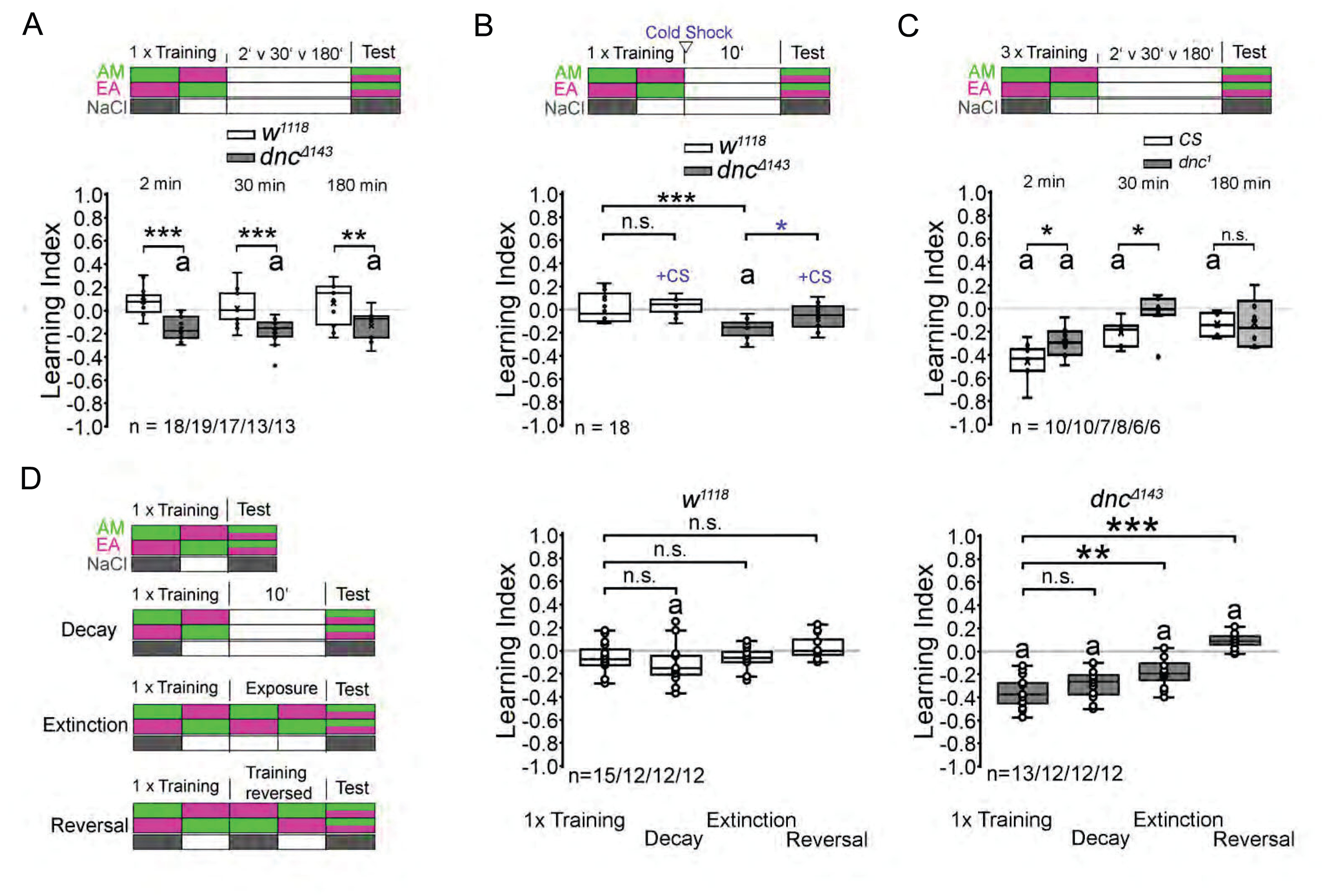
The improved learning and memory performance of *dnc*^Δ*143*^ mutants is long-lasting. ***A***, The memory formed after one training cycle in *dnc*^Δ*143*^ was observable at 2 min, 30 min and 180 min after training and differed significantly from that in the controls (2 min: *w^1118^* = -0.07 ± 0.03, *dnc*^Δ*143*^ = -0.17 ± 0.02, Df (35), t = 7.38, *P* = 1.21529E-08; 30 min: *w^1118^* = -0.01 ± 0.04, *dnc*^Δ*143*^ = -0.18 ± 0.03, Df (30), t = 4.32, *P* = 0.00016; 180 min: *w^1118^* = -0.05 ± 0.05; *dnc*^Δ*143*^ = -0.14 ± 0.03, Df (24), t = 3.26, *P* = 0.0033). ***B***, After one cycle of training, cold shock had no significant effect on the behavior of *w^1118^*control larvae (*w^1118^* = -0.01 ± 0.03; *w^1118^* + CS = -0.02 ± 0.02). However, cold shock immediately after training erased learning and memory performance in *dnc*^Δ*143*^mutants (*dnc*^Δ*143*^ = -0.16 ± 0.02; *dnc*^Δ*143*^ + CS = -0.06 ± 0.02; Df (16), t = -3.48; *P* = 0.0014). ***C*,** The memory formed after three training cycles in *dnc^1^* was observable 2 min after training but not 30 min or 180 min later (2 min: *CS* = -0.46 ± 0.05, *dnc^1^* = -0.3 ± 0.04, Df (19), t = -2.75, *P* = 0.014; 30 min: *CS* = -0.2 ± 0.04, *dnc^1^*= -0.03 ± 0.06, Df (14), t = -2.26, *P* = 0.024; 180 min: *CS* = -0.14 ± 0.04, *dnc^1^* = -0.13 ± 0.09, Df (11), t = -0.098, *P* = 0.925). ***A***, to ***C*,** Normally distributed data were analyzed with an unpaired Student’s t test. ***D***, After one training cycle, *w^1118^* larvae do not exhibit significant learning; there is also no memory decay, extinction, or reversal of learning (*w^1118^* = -0.06 ± 0.04; *w^1118^* decay = -0.11 ± 0.05; *w^1118^* extinction = -0.07 ± 0.03; *w^1118^* reversal = 0.03 ± 0.03; Df (3:47), F = 2.058, *P* = 0.119). After 10 min of retention, *dnc*^Δ*143*^ exhibits a memory. Presentation of the odorants during the delay did not significantly reduce learning or memory. The mutant significantly learned and remembered a new association between the odorants. *dnc*^Δ*143*^ = -0.36 ± 0.04; *dnc*^Δ*143*^ decay = -0.27 ± 0.05; *dnc*^Δ*143*^ extinction = -0.18 ± 0.04; *dnc*^Δ*143*^ reversal = 0.09 ± 0.02; Df (3:47), F = 19.621, *P* = 2.17E-05). ***D***, Normally distributed data were analyzed with one-way ANOVA and Tukey’s HSD test. Significant differences from random choice were determined for all the data using a one-sample t test and are indicated with the letter “a” (*P\** < 0.05; **: *P\*\** < 0.01; ***: *P\*\*\**< 0.001 and “n.s.” = nonsignificant).

Next, we investigated whether the enhanced memory of the *dnc*^Δ*143*^ mutants could be erased and whether memory affects the ability to learn new associations (Figure 2D). To analyze memory extinction, the larvae were first trained to associate an odorant with the reinforcer, followed by a second training session without the reinforcer. Because the training lasted 10 min longer than the normal training cycle, we first investigated whether the delay in testing affects memory performance. Although *w^1118^*did not show significant memory performance immediately after training, it did show significant memory performance after 10 minutes of retention time (Figure 2D). This suggests that a longer-lasting form of memory is formed after a training cycle in *w^1118^*. The *dnc*^Δ*143*^ mutants showed no significant memory loss within 10 min of training relative to larvae tested immediately after training (Figure 2D). Thus, the second exposure to odorants without the reinforcer did not result in significant memory extinction. The *dnc*^Δ*143*^ mutants learned to form a new association between the trained odorants immediately after a reversed training procedure. Thus, the *dnc*^Δ*143*^ mutants are “better learners”, and the learning and memory of new associations with the same odorants are not influenced by the strength of the previously formed memory.

### Different Dnc isoforms suppress the early onset of memory in *dnc^Δ143^* mutants

In *dnc^Δ143^* mutants, the *dnc-RA* and *dnc-RL* transcripts are downregulated (Ruppert et al., 2017). First, we addressed whether the reduction in *dnc* is indeed the cause of the improved learning and memory phenotype observed in the mutants. To this end, we expressed the RNA interference line *UAS*-*dnc ^KK108541^* (Fenckova et al., 2019) under the control of the *dnc^RA^*-Gal4 driver (Ruppert et al., 2017) and analyzed learning and memory (Figure 3A). After one cycle of training, the reduction in *dnc* resulted in improved memory. After one or three cycles of training, overexpression of the missing Dnc^PA^ isoform using a *UAS*-Dnc^PA^::GFP transgene under the control of the *dnc^RA^*-Gal4 driver did not improve or change learning or memory (Figure 3B and Figure 3C). These results are consistent with the hypothesis that the observed phenotype is caused by the loss of *dnc*.

**Figure 3:**
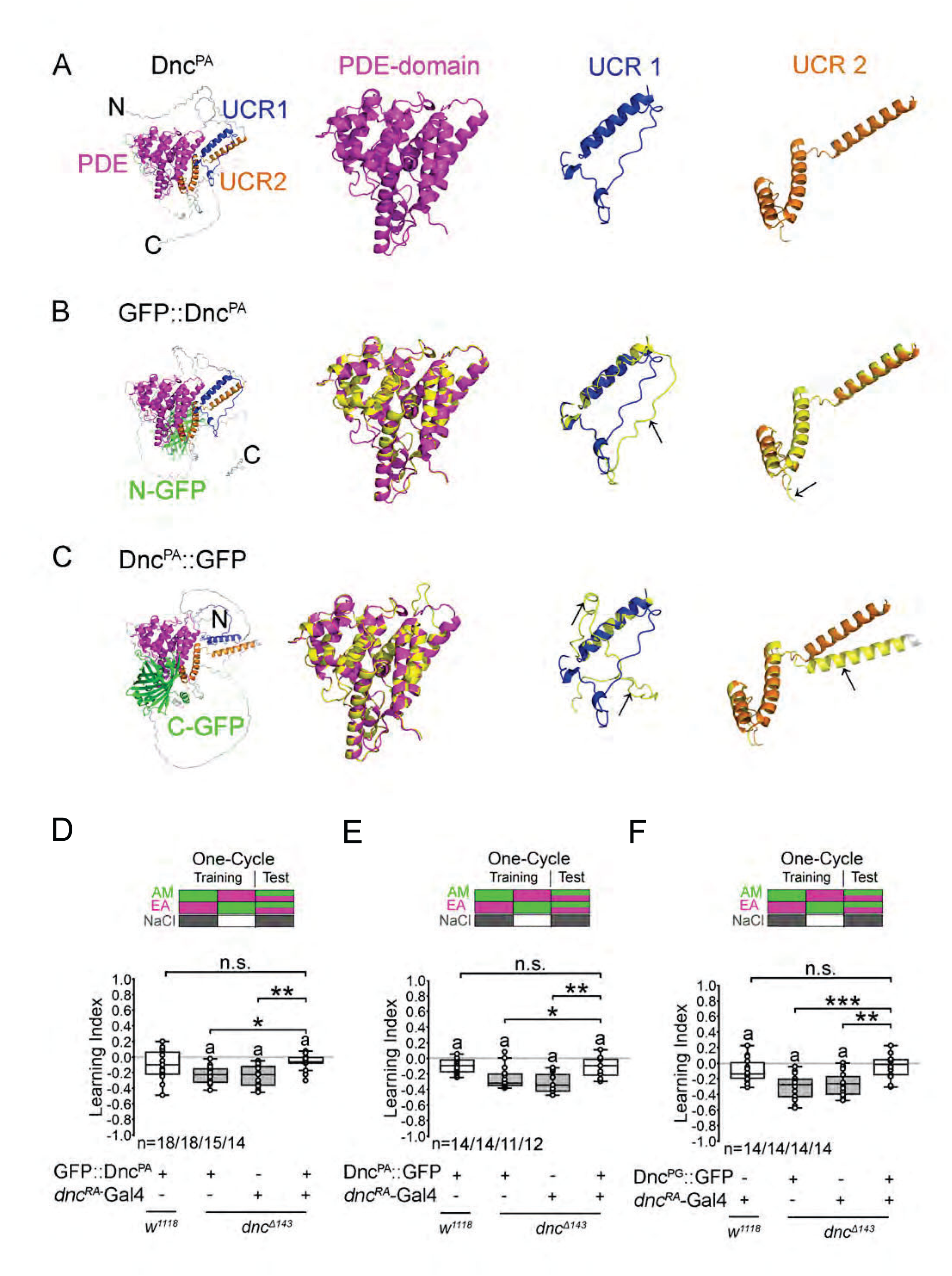
Two different Dnc isoforms repress the memory of *dnc^Δ143^*mutants. ***A-C*** The protein structures predicted using AlphaFold2 (Jumper et al., 2021) for ***A***, the endogenous Dnc^PA^ protein, ***B***, GFP::Dnc^PA^ and ***C***, Dnc^PA^::GFP were rendered using PyMOL2.6. The PDE domain of the endogenous proteins is shown in magenta; the UCR1 domain, in blue; and the UCR2 domain, in orange. The respective domains of the fusion proteins are marked in yellow, and the structural overlap is shown by the mixture of magenta, blue or orange with yellow. The arrows point toward structures that do not match. ***D***, The expression of GFP::Dnc^PA^ under the control of the *dnc^RA^-*Gal4 driver restored the learning and memory phenotype of the *dnc^Δ143^* mutant to control levels (Df (3,61), F= 6.813, P = 0.00041; UAS-GFP::Dnc^PA^= -0.09 ± 0.04; ; UAS-GFP::Dnc = -0.23 ± 0.03; *dnc* ; *dnc -*Gal4/+= -0.25 ± 0.04; *dnc*^Δ*143*^; UAS-GFP::Dnc^PA^; *dnc^RA^-*Gal4 = -0.07 ± 0.03). ***E***, The expression of Dnc^PA^::GFP restored learning and memory to control levels (Df (3,47), F = 8.405, P = 0.00014; UAS-Dnc^PA^::GFP = -0.1 ± 0.03; *dnc*^Δ*143*^; UAS-Dnc^PA^::GFP = -0.26 ± 0.04; *dnc*^Δ*143*^; *dnc^RA^-*Gal4/+= -0.31 ± 0.04; *dnc*^Δ*143*^; UAS-Dnc^PA^::GFP; *dnc^RA^-*Gal4 = -0.11 ± 0.04). ***F***, The expression of Dnc^PG^::GFP repressed early onset of memory in *dnc^Δ143^* mutants (Df (3,52); F= 9.374, P = 0.000047; *dnc^RA^-*Gal4/+ = -0.09 ± 0.04, *dnc*^Δ*143*^;UAS-Dnc^PG^::GFP = -0.31 ± 0.04, *dnc*^Δ*143*^; *dnc^RA^-*Gal4/+= -0.27 ± 0.04, *dnc*^Δ*143*^;UAS -Dnc^PG^::GFP; *dnc^RA^-*Gal4 = -0.05 ± 0.04). The normally distributed data from the behavioral experiments were analyzed using one-way ANOVA followed by Tukey’s HSD test. The letter “a” indicates significant differences from random choice as determined using a one-sample t test (P < 0.05). Differences between groups are marked as follows: *: P < 0.05, **: P < 0.01 and *** P < 0.001. For more details of the statistics, see Table S1.

Next, we investigated whether the expression of Dnc^PA^ could suppress the formation of memory in *dnc^Δ143^* mutants (Figure 4). In these experiments, we used two different *UAS-Dnc^PA^* transgenes that were either tagged with eGFP at the N-terminus or at the C-terminus of the protein (Ruppert et al., 2017). To address whether the GFP tag influences the structural integrity of the Dnc^PA^ protein, we used AlphaFold2 (Jumper et al., 2021) to simulate the folding properties of Dnc^PA^ and the tagged Dnc^PA^ proteins (Figure 4A to Figure 4C). The PDE domain was defined using UniProt (The UniProt Consortium, 2021), and the UCR domains were defined as previously described (Bolger et al., 1993). The GFP-tag on the N- or C-terminal region of Dnc^PA^ was predicted not to influence the structure of the PDE domain (Figure 4A to Figure 4C). The N-terminal region contains the UCR1 and UCR2 domains, in addition to Dnc isoform-specific sequences. The UCR domains are important for heterodimer and homodimer formation (Richter and Conti, 2002) (Xie et al., 2014). Differences in the structures of the UCR domains of the fusion proteins and Dnc^PA^ were observed depending on the site of the GFP tag (Figure 4B and Figure 4C). While the α-helical fragment of UCR1 overlaps with the region in Dnc^PA^ in both tagged proteins, the flanking regions do not overlap. The UCR2 domain of the N-terminally tagged Dnc^PA^ mainly overlaps with the UCR2 domain of the endogenous protein, although there is a slight shift in the regions that do not contain a known protein motif. In contrast, the ankle of one of the α-helices is strongly shifted in the UCR2 domains of the C-terminal tagged version of Dnc^PA^. Thus, the C-terminal-tagged Dnc^PA^ fusion protein differs more than the N-terminal-tagged Dnc^PA^. does from the endogenous Dnc^PA^ protein in its structural features. Interestingly, both fusion proteins repressed the early onset of memory in *dnc^Δ143^* mutants when expressed under the control of the *dnc^RA^*-Gal4 driver (Figure 4D and Figure 4E). Thus, the position of the tag at the N- or C-terminus of the protein did not interfere with the function of the PDE activity of the Dnc^PA^ protein, consistent with the prediction of no structural changes in the PDE domain.

**Figure 4:**
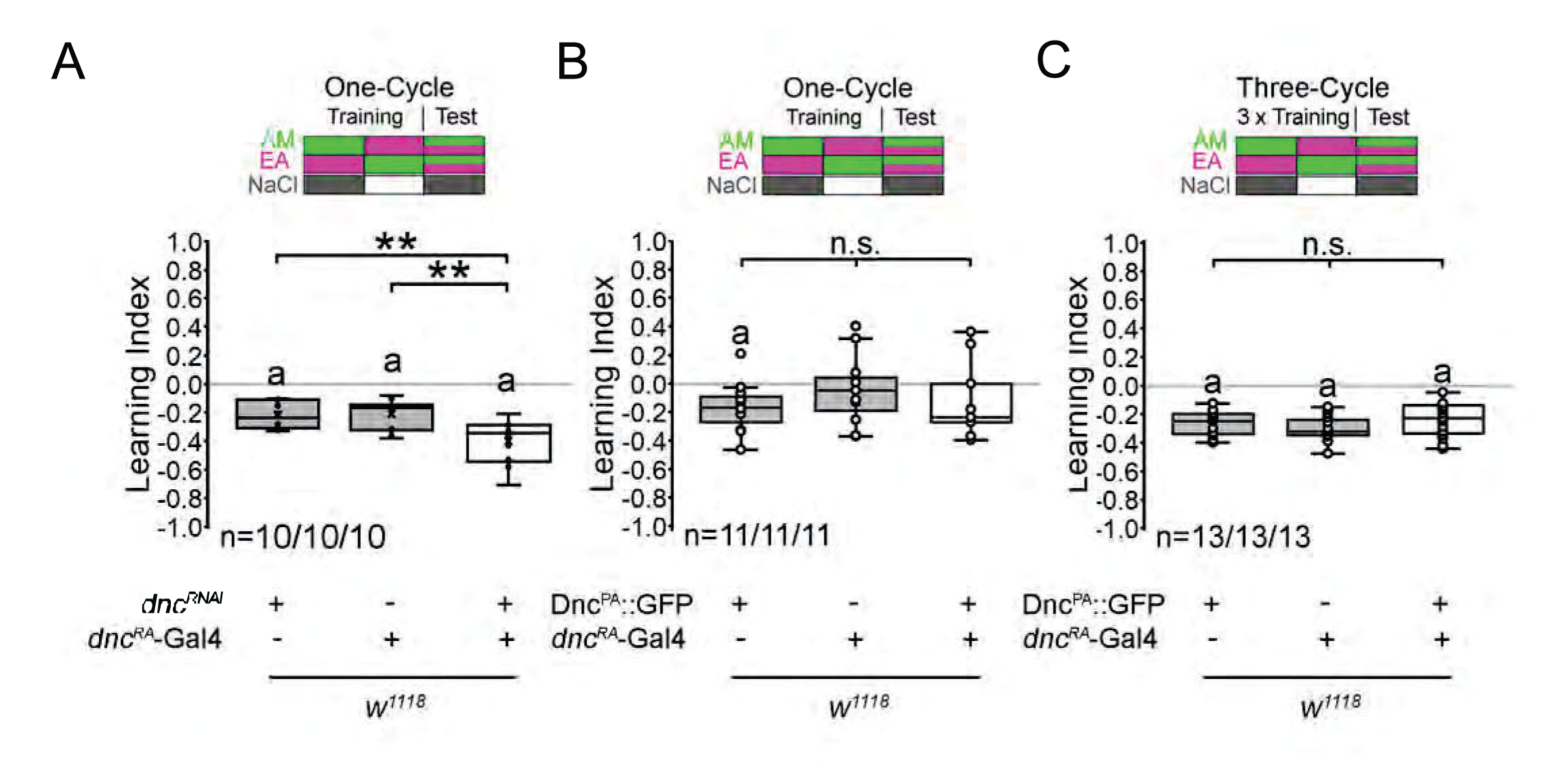
Reduced *dnc* expression improves memory. ***A*,** After one cycle of training, knockdown of *dnc* in *dnc^RA^*-positive cells significantly improved memory (Df (2,27); F= 8.181; P = 0.0017; UAS-*Dnc^RNAi^*= -0.21 ± 0.03; *dnc^RA^-*Gal4/+= -0.2 ± 0.03; *dnc^RA^-*Gal4; UAS-*Dnc^RNAi^* = -0.04 ± 0.05). **B-C,** The overexpression of Dnc^PA^ under the control of *dnc^RA^-*Gal4d did not significantly affect learning and/or memory in *w^1118^* larvae after **B,** one cycle of training (Df (2,30); F= 0.7999, P = 0.4587; UAS-Dnc^PA^::GFP = -0.17 ± 0.05, *dnc^RA^-*Gal4/+= -0.04 ± 0.07; *dnc^RA^-*Gal4; UAS-Dnc^PA^::GFP = -0.12 ± 0.08) or **C,** three cycles of training (Df (2,36); F= 1.4275, P = 0.2532; UAS-Dnc^PA^::GFP = -0.27 ± 0.02, *dnc^RA^-*Gal4/+= -0.3 ± 0.02, dncRA-Gal4; UAS-Dnc^PA^::GFP = -0.24 ± 0.03). The letter “a” indicates significant differences from random choice as determined using a one-sample t test (P < 0.05). Differences between groups were analyzed using ANOVA with Tukey’s post hoc test, with * indicating P < 0.05, ** indicating P < 0.01 and *** indicating P < 0.001. For details of the statistics, see Table S1.

To address whether other Dunce isoforms can substitute for the function of Dnc^PA^, we expressed the Dnc^PG^::GFP isoform under the control of the *dnc^RA^*-Gal4 driver in *dnc^Δ143^* mutants (Figure 4F). The Dnc^PG^ isoform differs from Dnc^PA^ in its N-terminal region, and it is the only isoform that contains a nuclear localization signal. The Dnc^PG^::GFP fusion protein was able to target the nucleus in follicle cells, which was not the case for Dnc^PA^::GFP (Ruppert et al., 2017). The expression of Dnc^PG^::GFP repressed the early onset of memory in *dnc^Δ143^* mutants. Thus, different Dnc isoforms are able to suppress the early onset of memory observed in *dnc^Δ143^* mutants.

### Subcellular localization of Dnc isoforms in neurons

To investigate where Dnc^PA^ function is required at the subcellular level, we analyzed the expression of Dnc^PA^ in neurons. Since no Dnc^PA^ isoform-specific antibody is available, we analyzed the expression patterns of the GFP fusion proteins (Figure 5). The GFP-tagged transgenes were targeted to the glutamatergic neurons at the neuromuscular junction (NMJ) using the *DvGlut*-Gal4 driver (Collins et al., 2006). For comparison, the expression of Dnc^PG^::GFP was also analyzed. As a reference, the membrane-targeted UAS-mCD8::GFP transgene was used. The Dnc^PA^::GFP and GFP::Dnc^PA^ fusion proteins are expressed in the soma but not in the nucleus. In contrast, Dnc^PG^::GFP is expressed in the soma and in the nucleus (Figure 5A).

**Figure 5:**
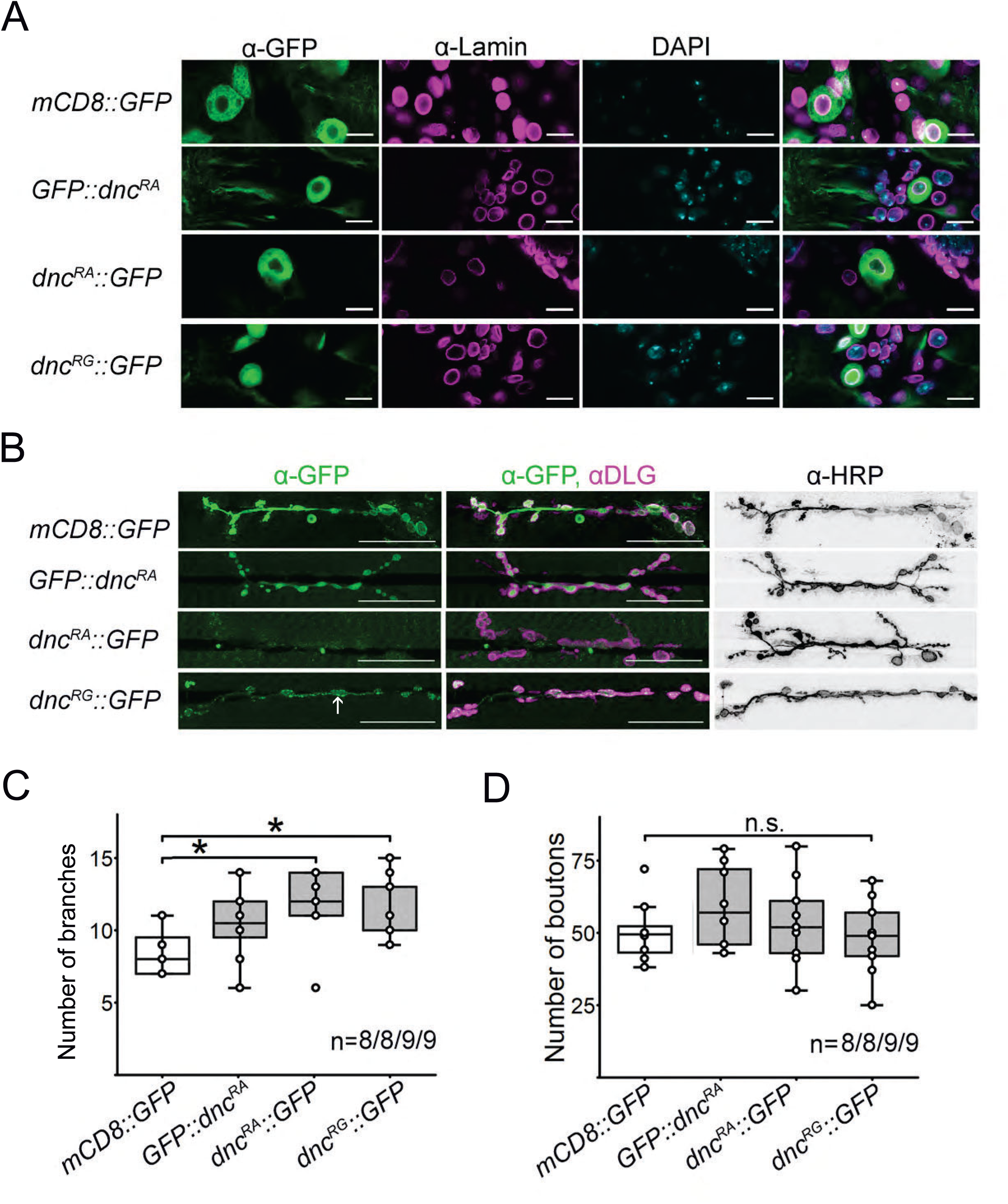
Subcellular localization of Dnc proteins. ***A*,** Dnc transgenes were expressed under the control of the *dvGlut*-Gal4 driver. The GFP fusion proteins are labeled in green. The nucleus was visualized using DAPI (blue), and the nuclear envelope was visualized with anti-Lamin antibodies (magenta). Dnc^PG^::GFP is expressed in the nucleus and the soma, whereas the Dnc^PA^ fusion protein is expressed only in the soma. The scale bar is 10 µm. ***B*,** The membrane of neurons at the NMJ was visualized using an anti-HRP antibody. GFP expression of GFP::Dnc^PA^ and Dnc^PG^, but not Dnc^PA^::GFP, was detected at NMJ 6/7 in A4. The scale bar is 50 µm. ***C*,** The number of branches was significantly increased in larvae expressing Dnc^PA^::GFP and Dnc^PG^::GFP. ***D***, There was no significant difference in the number of boutons at the NMJ in the larvae overexpressing the isoforms. For ***C*** and ***D*,** the number below the box indicates the number of NMJs analyzed. Differences were analyzed using ANOVA with Tukey’s post hoc test. *: P < 0.05, **: P < 0.01 and *** P < 0.001.

At the axon terminals, the N-terminal tagged Dnc^PA^ was restricted to the membrane of the synaptic varicosities. At the varicosities, the expression resembled that of the membrane-bound GFP (for comparison, see Figure 5B panels 1 and 2). C-terminal-tagged Dnc^PA^ was not expressed at the NMJ. C-terminal-tagged Dnc^PG^ appeared in a dot-like pattern within the synaptic varicosities (Figure 5B).

The overexpression of the PDE domain of Dnc reduced the number of synaptic varicosities (Cheung et al., 1999). Although the expression of the Dnc fusion proteins appeared to change the morphology at the NMJ (Figure 5B). To quantify these possible changes, we counted the number of branches and the number of synaptic varicosities (Figure 5C and Figure 5D). The overexpression of Dnc^PA^::GFP and Dnc^PG^::GFP increased the number of branches significantly but did not change the number of synaptic varicosities. The overexpression of GFP::Dnc^PA^ did not change the number of branches or varicosities; however, the variance in the number of branches was quite high. Thus, similar to the overexpression of the PDE domain, the overexpression of Dnc^PA^ and Dnc^PG^ alters neuronal morphology; however, the changes differ. While overexpression of PDE decreased in the number of varicosities, overexpression of the fusion protein increased the numbers of branches. The presence of the N-terminal regions of Dnc proteins appears to influence the types of changes in neuron morphology.

In summary, the expression of GFP::Dnc^PA^, Dnc^PA^::GFP and Dnc^PG^::GFP in the soma is similar, but their expression at the NMJ differs. The overlapping expression pattern of the fusion proteins suggests that the function of Dunce in the soma of neurons is required to suppress the early onset of memory.

### The *dnc^RA^-*Gal4 driver targets a small group of neurons

To identify where in the CNS Dunce is required to suppress memory in *dnc*^Δ*143*^ mutants, we visualized the Gal4 expression domain of the *dnc^RA^-*Gal4 driver using a UAS-mCD8::GFP transgene (Figure 6). In the brain and the ventral nerve cord (VNC), approximately 76 neurons in 21 clusters expressed GFP (Figure 6A and Figure 6B). In addition, the SStrhl cluster, a prominent cluster consisting of many cells in the optic lobes in the central brain, was also GFP positive.

**Figure 6:**
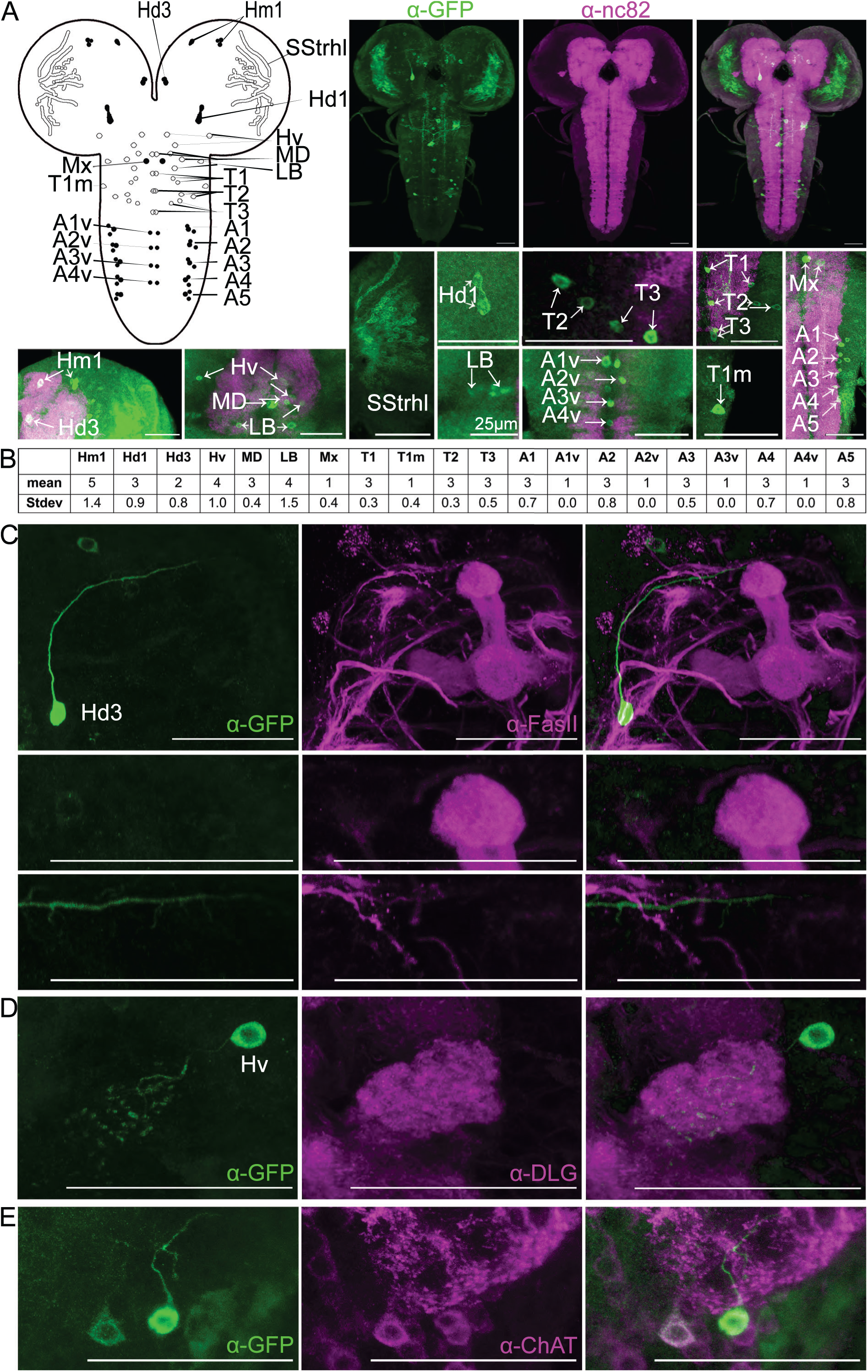
The expression domain of *dnc^RA^-*Gal4 in the larval CNS. ***A***, *dnc^RA^-*Gal4 drives the expression of UAS*-*mCD8::GFP transgenes in the larval CNS. The synaptic marker α-nc82 (magenta) was used as a reference for the brain regions. The cartoon illustrates the identified neurons, and Table B summarizes the identified clusters. At least six larval CNSs were analyzed. ***C*, *D***, UAS-tau::GFP transgene was used to visualize the projection of the *dnc^RA^*-Gal4 driver-targeted neurons. ***C*,** The mushroom bodies were labeled using an α-FasII antibody (magenta). One neuron of the Hm1 cluster projects near the mushroom bodies but does not enter the structure. ***D***, The larval antennal lobes were visualized using an antibody against α-Disc-large (DLG; magenta). One neuron of the Hv-cluster projects into the antennal lobe. ***D***, Two Hv-neuronal neurons are positive for ChAT. Scale bars are 50 µm unless otherwise indicated.

The simple model of how associative olfactory learning and memory form in the larval brain suggests that coincidence detection is necessary for the association between conditioned and unconditioned stimuli in larval mushroom room bodies and leads to increased cAMP levels. The odor information is transferred via cholinergic neurons onto the mushroom bodies and the reinforcer by dopaminergic neurons (Widmann et al., 2018). Here, four types of dopaminergic neurons that project to larval mushroom bodies mediate punishment in the larval brain (Weber et al., 2025). Therefore, we next asked whether neurons that require Dunce are part of the mushroom bodies or form contacts with the mushroom bodies.

The soma of one neuron of the Hd3 cluster is located near the mushroom bodies, but the neuron does not project into the mushroom bodies (Figure 6C). In addition, the neurons of the Hm1 cluster, Hd1 cluster, MD cluster, and LB and Mx clusters do not project into the vicinity of brain regions involved in olfactory information processing, such as the larval antennal lobes (LAL) and mushroom bodies. Only one neuron of the Hv cluster projected into the glomeruli of the LAL (Figure 6D). Two additional neurons of the cluster project into the region below the antennal lobe, the ventromedial cerebrum (VMC; Figure 6E). These neurons express choline acetyl transferase (ChAT), a marker of cholinergic neurons (Figure 6E). In summary, we did not detect any evidence that Dunce is required in mushroom bodies to suppress the early onset of memory.

## Discussion

We determined that the *dnc*^Δ*143*^ mutants learned and remembered better than the known *dnc^1^*mutants did. cAMP levels are increased in *dnc*^Δ*143*^ mutants (Ruppert et al., 2017). The phenotype of the *dnc*^Δ*143*^ mutants and the increased cAMP levels are in line with the expectation that elevated cAMP should increase neuronal activity and thereby strengthen the synapses required for learning and memory. Adding PDE activity back to the mutant reduced the improvement in learning and memory performance. We pinpointed the requirement for phosphodiesterase activity to a subset of neurons, none of which innervate the mushroom bodies, a structure that is thought to mediate the coincident detection of the reinforcer and the conditioned stimulus by elevating cAMP levels. Our comparative expression studies of different Dunce isoforms further suggested that at the subcellular level, PDE activity is required in the somatic region of neurons, a region that is involved in the integration of synaptic input to the generation of the action potential and firing rate of a neuron.

### What kind of memory is affected in *dnc***^Δ^***^143^* mutants?

Classic *dnc^1^* mutant larvae exhibit defects in short-term memory that become clearly visible after three rounds of training (Khurana et al., 2009). This deficiency has been attributed to a failure to form short-term memory, specifically anesthesia-sensitive short-term memory (Widmann et al., 2016). The *dnc*^Δ*143*^ mutant does not affect the same form of memory.

First, there are characteristic differences: *dnc*^Δ*143*^ shows the opposite phenotype, with enhanced memory after less training, and this memory is still measurable after three hours, which is not the case for *dnc^1^*mutants. The opposing phenotypes of *dnc^1^* and *dnc*^Δ*143*^ could be due to differences in Dnc expression, e.g., loss versus elevation of Dnc expression. In *dnc^1^* mutants, most *dunce* transcripts are downregulated, including the *dnc^RA^* transcript that is missing in *dnc*^Δ*143*^ (Ruppert et al., 2017). Our work clearly revealed that the phenotype of the *dnc*^Δ*143*^ mutant is due to the loss of Dnc^PA,^ as the *dnc*^Δ*143*^ mutant is a *dnc^RA^-*specific mutant; restoring PDE activity and Dnc^PA^ expression in the mutant repressed learning to control levels, and overexpression did not change the STM or LTM in the neurons required to regulate this behavior. In *dnc^1^* mutants, the *dnc^RA^* transcript level is reduced, but not to the same extent. If reducing the *dnc^RA^* transcript were sufficient to improve learning and memory, both mutants should share the same phenotype, but this is not the case. Furthermore, the loss of *dnc^RA^* does not account for defects in memory after more training, since there are no differences in memory between control and *dnc*^Δ*143*^ mutants after multiple training sessions. Consistent with this idea, the reduction in Dnc results in improved memory because in adult flies, knockdown of *dnc* in serotonergic neurons also results in increased LTM (Scheunemann et al., 2018). Taken together, these results are consistent with the hypothesis that the defects in *dnc^1^* are due to the dysregulation of Dunce isoforms other than *dnc^RA^*.

Second, different neurons regulate the increases and reductions in memory. PDE activity under the control of the *dnc^RA^*-Gal4 driver suppressed learning and memory in *dnc*^Δ*143*^ mutants but did not restore the loss of memory in *dnc^1^*. In addition, overexpression of PDE did not result in decreased memory after three cycles of learning, when the memory defect of *dnc^1^* became clearly visible. The *dnc*^Δ*143*^ and *dnc*^1^ mutants have elevated cAMP levels (Ruppert et al., 2017; Davis and Kiger, 1981), but the phenotypes indicate opposing functions. At the network level, this might be achieved by Dnc-dependent regulation of cAMP in excitatory or inhibitory neurons. Together, they might be able to regulate the gain and loss of memory.

Another form of learning and memory that might be affected in the *dnc*^Δ*143*^ mutant is habituation. A failure to habituate might result in improved memory shortly after one cycle of training. Dnc knockdown results in loss of odorant habituation (Fenckova et al., 2019). However, habituation is not as long-lasting as the observed memory in *dnc*^Δ*143*^ mutants. Given that the memory is sensitive to cold-shock, is long-lasting and differs phenotypically from *dnc^1^*, Dnc^PA^ is required to suppress the early onset of this long-term memory. At the subcellular level, the phenotypic analysis of the Dnc fusions suggests that cAMP needs to be tightly regulated in the soma to suppress long-term memory.

### What kind of neurons inhibit memory?

The neurons in which PDE activity is required might provide insight into the kind of memory that is enhanced in *dnc*^Δ*143*^. In the larval brain, as in the adult brain, the mushroom bodies are considered the ‘memory’ center, where the information of the conditioned stimulus and the odor mediated by cholinergic projection converge with the information of the unconditioned stimulus mediated by biogenic amine-expressing neurons to form memory (Widmann et al., 2018). Increased cAMP signaling is considered a readout for memory formation (Gervasi et al., 2010). A reduction in the activity zone protein Bruchpilot in larval mushroom bodies results in a reduction in short-term memory and the elimination of memory that emerges after multiple training cycles when salt is used as a negative reinforcer (Widmann et al., 2016). In addition to these forms of memory, mushroom bodies also regulate habituation, a nonassociative form of learning. For example, a reduction in the activity of the choline transporter in larval mushroom bodies results in a loss of habituation and hypersensitivity to the odor (Hamid et al., 2021). In the adult fly brain, a reduction in Dnc in the serotonergic neurons engulfing mushroom bodies facilitates long-term memory (Scheunemann et al., 2018). However, we did not find evidence that Dnc^PA^ or PDE activity is required in neurons that are intrinsic to mushroom bodies or in neurons that project into or surround the mushroom bodies. These findings suggest that neurons outside the mushroom body suppress long-lasting memory.

The *dnc^RA^*-Gal4 driver targets one neuron that innervates the larval antennal lobes and neurons along with cholinergic neurons that innervate the ventromedial cerebrum and both structures of the deuterocerebrum and neurons that innervate the mandibular neuromeres. These regions process input from the olfactory and gustatory sensory system and thus process the information associated with the conditioned and unconditioned stimulus (Kendroud et al., 2018; Python and Stocker, 2002). We did not observe any changes in odorant attractiveness in the mutants or controls, nor did we observe that the reinforcer changed the perception of the odorants. However, it is possible that the improvement in memory is due to a prolongation of the stimulation by the conditioned stimulus or unconditioned stimulus, which needs to be further investigated.

Memory inhibition occurs in neurons that neither directly innervate nor compose any part of the mushroom bodies, suggesting that Dunce regulates learning and memory in neurons that modulate the sensory input or reward, similar to what has been observed in other animal models of learning and memory, such as Aplysia. Comparison of the cellular localization of different Dunce isoforms revealed that the regulation of cAMP in the soma might be crucial to suppress early onset of memory. Thus, depending on the neuron type, elevated cAMP levels increase or reduce memory.

## Acknowledgments

We thank the members of the Scholz laboratory for their continued discussions and Victoria Brüning, Evelin Fahle and Anniwaer Kaiweisaer for data collection. Stocks obtained from the Bloomington *Drosophila* Stock Center were used for this study. The work was supported by the Deutsche Forschungsgemeinschaft DFG. To H.S. DFG656 SCHO10-1.

## Author contributions

TH and HS designed the research; TH, MV, AK, VB, MM, and MG performed the research; TH and HS analyzed the data; and TH and HS wrote the paper. Funding acquisition: HS acquired funding.

## Supplemental material

The supplementary material for this paper includes **Figure S1,** related to Figure 1, and **Table S1,** containing data and statistics for the behavioral experiments.

**Figure S1 Learning and memory of *dnc*^Δ*143*^ mutants when AM and BA are used as odorants *A***, After one training session, control animals showed memory when AM and BA were used as odorants, whereas *dnc^1^* mutants did not. The *dnc*^Δ*143*^ larvae learned to associate the aversive cue of 2 M NaCl with the odorant more effectively than did the control animals. *CS* = -0.18 ± 0.4, *dnc^1^* = -0.03 ± 0.04, Df (14), t = -3.136, *P* = 0.0073 and *w^1118^* = -0.12 ± 0.03*, w^1118^dnc*^Δ*143*^ = -0.26 ± 0.04, Df (27), t = -2.953, *P* = 0.0064. ***C***, After three training cycles, the *dnc^1^* and *dnc*^Δ*143*^ larvae learned and remembered as well as the larvae in their respective control groups. *CS* = -0.2 ± 0.03, *dnc^1^* = -0.14 ± 0.04, Df (32), t = -1.325, *P* = 0.1947 and *w^1118^* = -0.25 ± 0.06*, w^1118^dnc*^Δ*143*^ = -0.36 ± 0.04, Df (25), t = -1.483, *P* = 0.1505. Normally distributed data were analyzed with an unpaired Student’s t test. Significant differences from random choice were determined using a one-sample t test (*P* < 0.05) and labeled “a”. “n” indicates the number of reciprocally trained independent groups of larvae.

**Table S1: Data and statistics.**

