## Supplementary figures and images for "The PDE4D ortholog Dunce suppresses memory in *Drosophila melanogaster*"

### Supplement Figure 1

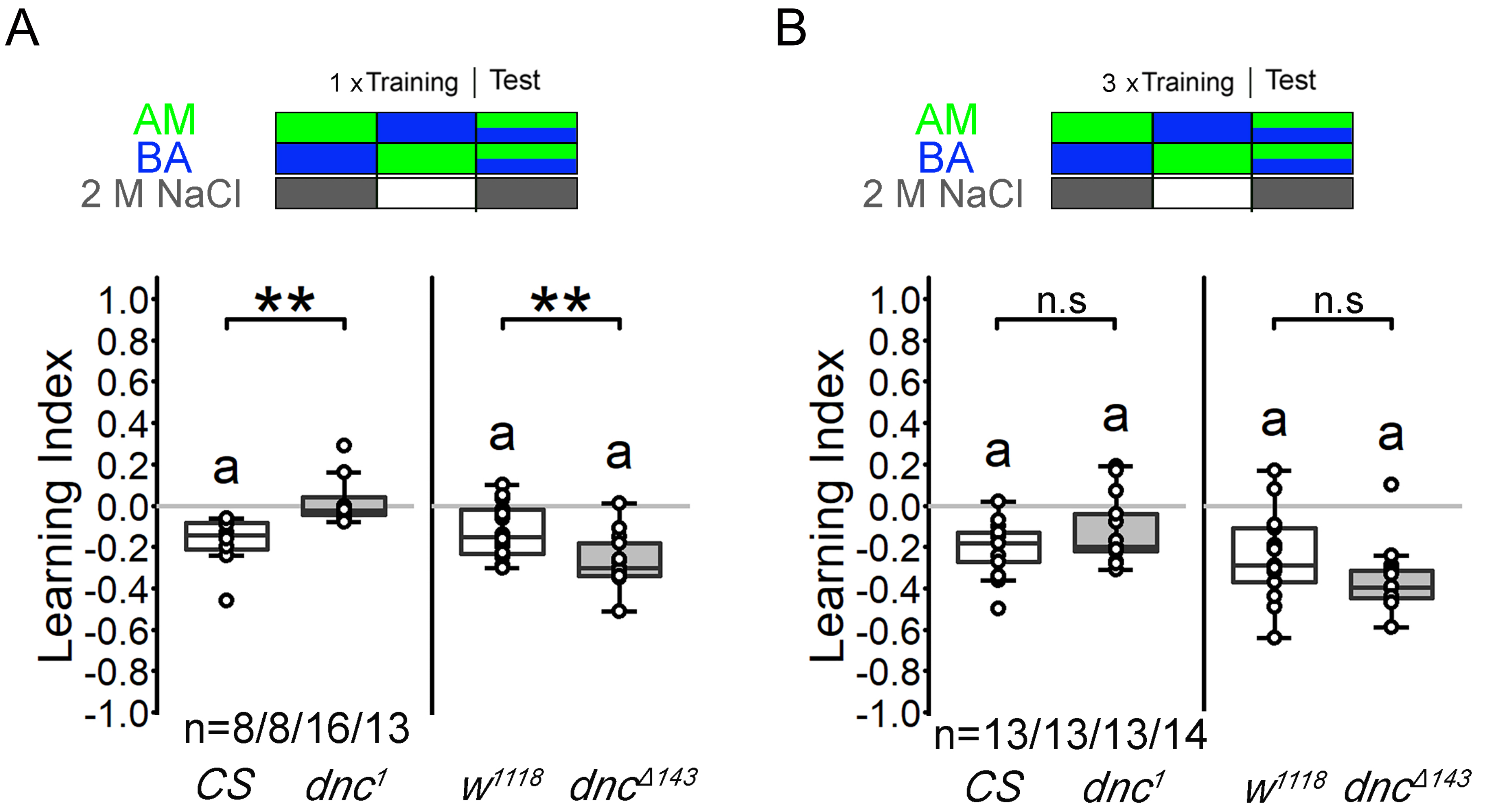
